# Calcium-driven In Silico Inactivation of a Human Olfactory Receptor

**DOI:** 10.1101/2024.01.31.578070

**Authors:** Lorenza Pirona, Federico Ballabio, Mercedes Alfonso-Prieto, Riccardo Capelli

**Affiliations:** Department of Biosciences, Università degli Studi di Milano, Via Celoria 26, I-20133 Milano, Italy; Computational Biomedicine, Institute for Neuroscience and Medicine INM-9, Forschungszentrum Jülich GmbH, Wilhelm-Johnen-Straße, D-54248 Jülich, Germany

## Abstract

Conformational changes as well as molecular determinants related to the activation and inactivation of olfactory receptors are still poorly understood due to the intrinsic difficulties in the structural determination of this GPCR family. Here, we perform, for the first time, the *in silico* inactivation of the human olfactory receptor OR51E2, highlighting the possible role of calcium in this receptor state transition. Using molecular dynamics simulations, we show that a divalent ion in the ion binding site, coordinated by two acidic residues at positions 2.50 and 3.39 conserved across most ORs, stabilizes the receptor in its inactive state. In contrast, protonation of the same two acidic residues is not sufficient to drive inactivation within the *`µ`*s timescale of our simulations. Our findings suggest a novel molecular mechanism for OR inactivation, potentially guiding experimental validation and offering insights into the possible broader role of divalent ions in GPCR signaling.

---

Olfactory receptors (ORs) are a family of G protein-coupled receptors (GPCRs) implicated in odor perception.^1^ This group of class A GPCRs comprises approximately 400 members, representing half of the GPCRs encoded in the human genome.^2^ The function of ORs is not limited to olfaction, as they have been identified in various extranasal tissues and have been shown to play a role in a plethora of physiological and pathological processes.^3,4^ This important clinical perspective, combined with the fact that non-olfactory GPCRs are the target of 34% of the total number of FDA-approved drugs,^5^ makes the study of the OR family one of the most promising fields for biochemistry and pharmacology.

Despite their potential as drug targets, ^6^ the study of ORs has been hindered by the lack of structural data: the first experimental structure was published in March 2023,^7^ consisting of human OR51E2 in its active state, bound to an agonist (propionate) and a mini-G protein. In light of this limitation, the modeling community has approached the OR family with different techniques, from homology modeling^8–11^ to *de novo* structure prediction.^12,13^ Such computational models have provided useful insights into the mechanism of OR-ligand recognition at the atomistic level, as well as pinpointed crucial residues for OR function, in combination with experimental mutagenesis data.

Recent cryoEM work by Choi *et al.*^14^ unveiled, for the first time, both the active and inactive conformations of a chimeric OR, OR52_cs_, which was constructed using a consensus sequence strategy based on 26 members of the OR52 family. The availability of experimental structural data for both conformational states of the same OR receptor opens the opportunity for cross-validating *in silico* models. Furthermore, this enables a computational exploration of the activation or inactivation process, aligning with the approach taken in similar computational studies on other class A GPCRs.^15,16^

In this study, we focus on the inactivation of human OR51E2, leveraging the cryoEM structure of this receptor in its active state.^7^ Our primary objective is to simulate the inactivation of the receptor *in silico* over a *µ*s timescale, drawing upon the pioneering work of Dror *et al.* on the *β*_2_-adrenergic receptor.^15^ Moreover, OR51E2, along with 90% of the OR family members, bears two acidic residues in the ion binding site^13^ (2.50 and 3.39 using the Ballesteros-Weinstein class A GPCR generic numbering^17^), indicating the potential involvement of these charged residues in the ion binding mechanism and conformational dynamics of ORs. This hypothesis is supported by our previous observations of sodium-bound OR51E2 in the inactive state.^13^ Accordingly, we aim at elucidating the role of the ion binding site for OR conformational plasticity, also building on previous work on sodium binding to other non-olfactory class A GPCRs.^18,19^

## Simulations with no ions in the binding site

Initially, the receptor structure was built from its experimentally determined active conformation (PDB code: 8F76^7^), with the agonist and mini-G protein removed. We employed the CHARMM-GUI web server^20,21^ for receptor set up and its embedding into a 3:1 POPC:Cholesterol bilayer (see Methods section and Supporting Information for further details). Following system preparation, we adopted a multi-step equilibration protocol, applying restraints to preserve the initial fold while relaxing the system (refer to Methods), as done in our previous work. ^13^ This was followed by five independent unrestrained MD simulations, each lasting 5 *µ*s.

In all five apo OR51E2 replicas, a partial loss of the receptor fold was consistently observed (see Figure 1a), which is in line with the results obtained when starting from *de novo* structures of OR51E2.^13^ The region most impacted by this partial unfolding was the interface between transmembranes helices 6 and 7 (TM6 and TM7, respectively), which widens significantly during the simulation (from 8 Å to 15 Å).

**Figure 1:**
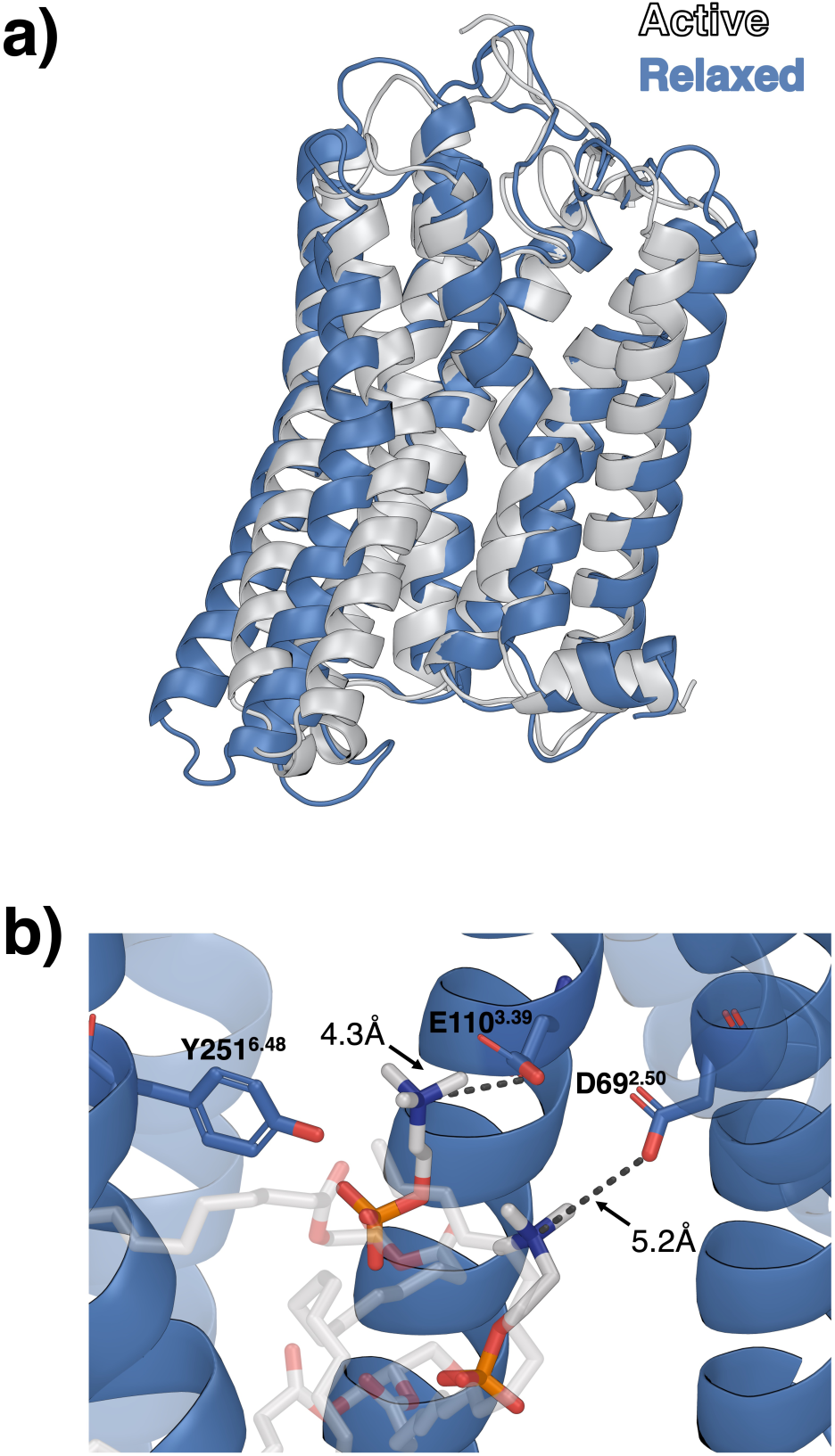
OR51E2 from simulations without ions in the binding pocket. (a) Superposition of the cluster centroid obtained from the five trajectories (blue) started from the cryoEM structure in its active state (white), evidencing the widening of the TM6-TM7 interface. (b) Detail of the ion binding pocket, highlighting the interaction between the charged residues D69^2.50^ and E110^3.39^ and the two POPC headgroups entering the receptor bundle. Additionally, Y251^6.48^ is oriented towards the ion binding pocket. Hydrogen atoms have been omitted for clarity, and the distances refer to the frame shown.

Additionally, the expansion of the TM6-TM7 interface coincides with POPC molecules “diving” into the ion binding pocket, as their positively charged choline groups orient towards the charged residues D69^2.50^ and E110^3.39^ (see Figure 1b). These events clearly indicate a significant charge imbalance near the ion binding pocket, leading to lipid snorkeling through the apolar membrane. Within this structural rearrangement, we observed several phenomena related to the system conformational dynamics: (i) Water molecules freely traversed from the intracellular to the extracellular side of the receptor; (ii) The toggle switch Y251^6.48^ es-tablished multiple transient interactions with the charged headgroups of POPC lipids (both choline and phosphate moieties), E110^3.39^, and S111^3.40^; and (iii) The negatively charged side chains of D69^2.50^ and E110^3.39^, whose charge is locally unbalanced, interacted with nearby polar and neutral amino acid side chains, and also electrostatically attracted the charged groups that entered the ion binding site (see Table S2 in the Supporting Information). Remarkably, we observed spontaneous binding of a Na^+^ ion in two out of the five simulations (see Table S2 and Figure S1). The ion entered from either the extra- (replica 4) or intracellular (replica 1) side of the membrane; however, it is important to note that, due to the use of periodic boundary conditions (PBC), the ionic concentration is nearly identical on both sides of the membrane.

As an independent confirmation of our observation of an electrostatic interaction between the ion binding pocket and charged groups, we considered the final frame of 1 *µ*s simulation started from the apo structure of OR52_cs_ (retrieved at https://github.com/sek24/natcomm2023); such simulation was performed under similar conditions to ours (except for the use of KCl instead of NaCl). Also in the case of OR52_cs_, two POPC molecules snorkel into the ion binding pocket. However, in our simulations the two lipid molecules wedge themselves between TM6 and TM7, whilst in the simulations of Choi *et al.* they do between TM5-TM6. In addition, the final OR52_cs_ frame shows a positively charged ion (K^+^) located close to D69^2.50^ and E110^3.39^, similar to the spontaneous sodium binding event observed in two of our replica simulations.

## Simulations with sodium in the ion binding site

After observing the spontaneous binding of Na^+^ in the ion binding site, we carried out a new set of five replica simulations. In these simulations, Na^+^ was manually positioned in the ion binding pocket of the initial structure, while retaining the same procedure and parameters as in the previous section. The simulation length was 5 *µ*s per replica. This approach aligns with the observations from our previous work on the *de novo* structures of the same receptor in the inactive state,^13^ which maintained their fold only when sodium was present in the ion binding pocket. However, here we began with the cryoEM structure of OR51E2 in the active state^7^ and conducted more and longer replicas.

Overall, the simulations involving Na^+^ showed slightly enhanced receptor stability (see Figure 2), which is reflected in less POPC snorkeling into the ion binding pocket (Figure S4), reduced water permeability (Table S2), and lower RMSD values (Figure S3) of the C*_↵_* atoms of the transmembrane domains with respect to the cryoEM structure. Differently from the simulations with no ions, two trajectories (replicas 1 and 3) did not reveal any POPC snorkeling. In the other three trajectories, we observed the POPC head group moving towards the ion binding site, passing either through the interface between TM5-TM6 and interacting mainly with E110^3.39^ (replica 5 and the last 50 ns of replica 2) or between TM6-TM7, engaging residues D69^2.50^ and E110^3.39^ (replicas 4 and 5). Interestingly, Na^+^ unbinding was detected in replica 5 in the last *µ*s, where the ion moved to the intracellular space. Shortly after this event, a second POPC entered the ion binding site. Finally, a new Na^+^ ion spontaneously translocated from the extracellular space to the ion binding site within 1 *µ*s from the initial dissociation event (Figure S3 and Table S2). This phenomenon supports, once again, the need for a charged group at this location, counteracting the negative charge of the two acidic residues at positions 2.50 and 3.39.

**Figure 2:**
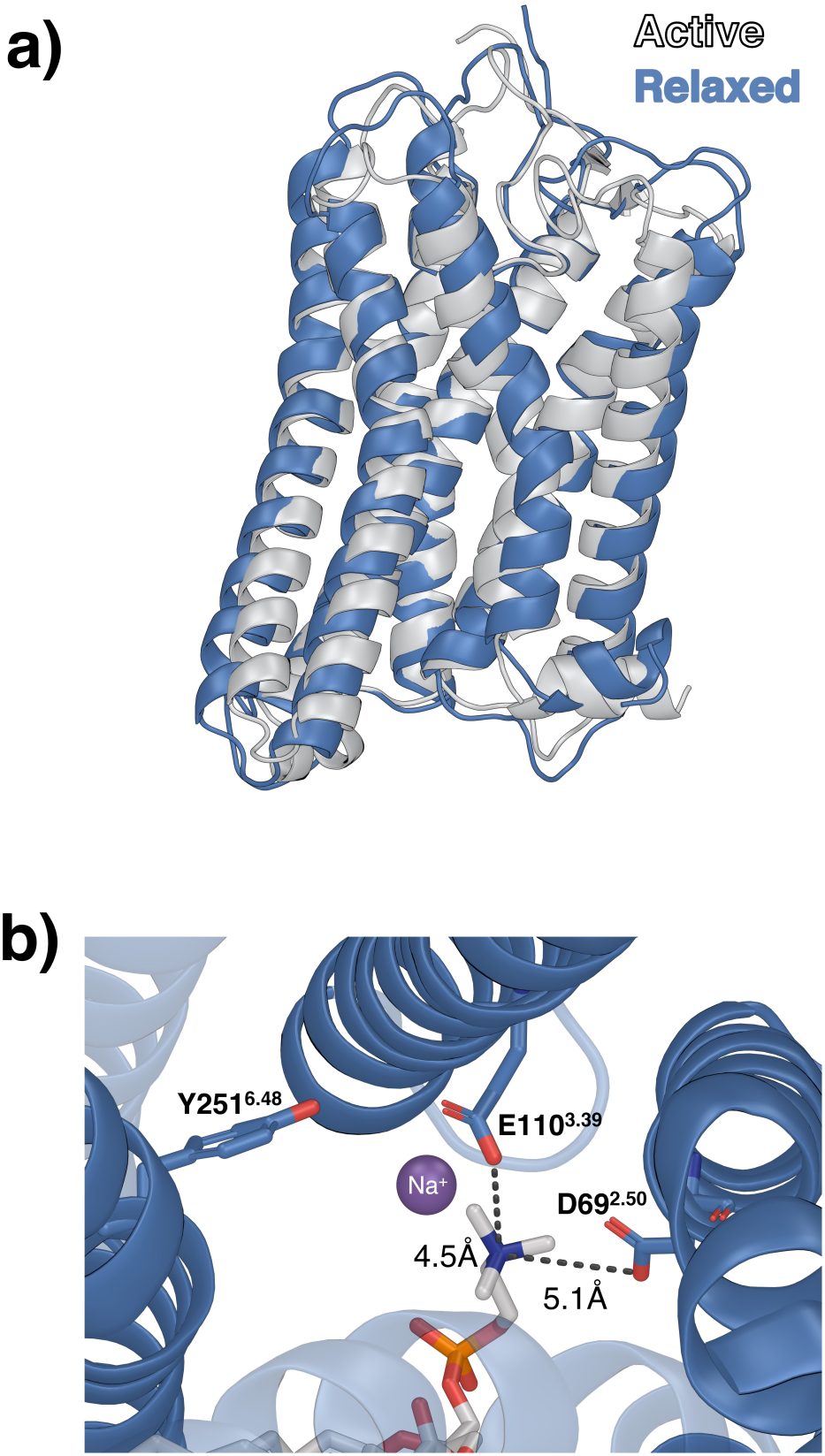
OR51E2 from simulations with Na^+^ ion in the binding pocket. (a) Superposition of the cluster centroid obtained from the five trajectories (blue) on the cryoEM structure in its active state (white). (b) Detail of the ion binding pocket, highlighting the interactions between Na^+^ and D69^2.50^ and E110^3.39^, as well as between POPC and the same two acidic residues. Hydrogen atoms have been omitted for clarity, and the distances refer to the frame shown.

In terms of water permeability, we observed reduced water passing through the protein compared to the simulations with no ions (Table S2). This trend was evident in the first three replicas, particularly in the second trajectory. Considering that the formation of a hydrated pathway spanning the receptor is connected to the reorganization of the H-bonds around the ion binding site, ^19^ we focused on replica 2 to identify changes in non-bonded interactions at the atomistic level. Namely, we performed a comparative H-bond analysis across all the ten replicas described thus far (five with Na^+^ and five without) to identify H-bonds that, in terms of occupancy, exhibited the most significant differences between the second replica and the others. The interaction between D69^2.50^ and Y291^7.53^ was particularly notable. The total occupancy of this H-bond (i.e. considering both oxygen atoms of the side chain of D69^2.50^) in the second replica of the Na^+^-bound system persisted for 76% of the trajectory. In contrast, replicas 1, 3, 4, and 5, showed occupancies of 19%, 52%, 3%, and 4%, respectively. Extending this analysis to the replicas without the ion in the binding site, this occupancy reached a maximum of 6% (replica 4), while in all the others it was around 1%. Y291^7.53^ has been reported as a conserved residue in class A GPCRs, implicated in the diffusion of Na^+^ ions,^18,19^ the formation of a water pathway inside the receptor, and in the receptor activation mechanism. ^22^ Another interesting H-bond pattern was observed between S111^3.40^ and Y251^6.48^, which is significantly more frequent in the presence of sodium. This increased persistence is particularly evident in three out of five trajectories with sodium (occupancy of 41%, 78%, 92%, 78%, 4% in replicas 1-5, respectively). In contrast, only one replica from the trajectories lacking ions exhibits a stable H-bond between these two residues (occupancy of 90% in replica 1, 0% in all the others).

To assess the stability of the receptor fold, we performed an RMSD calculation using the cryoEM structure of OR51E2 in its active state^7^ as the reference. Overall, the simulations with Na^+^ bound maintained the fold better (i.e. lower RMSD values - see Figure S3) than the simulations without ions (Figure S1), with some minor variability among them. Interestingly, peaks in the RMSD time evolution are linked to lipid snorkeling and increased water permeability in the receptor (Figure S4).

## Simulations with calcium in the ion binding site

From the previous simulations we can conclude that the charge imbalance in the ion binding site is still present, even if attenuated, when Na^+^ is bound. Hence, we decided to perform another round of five independent simulations, positioning a Ca^2+^ ion in the binding site while maintaining all the simulation parameters previously used (see Methods and Supporting Information). From a technical point of view, to minimize the limitations of the point charge representation of divalent ions, we manually placed calcium in the ion binding site, rather than trying to simulate its diffusion from the bulk solvent. Based on sequence analyses, class A GPCRs containing two acidic residues in the ion binding pocket have been hypothesized to bind Ca^2+^;^23^ however, to the best of our knowledge, this possibility has not yet been investigated in the context of ORs.

The simulations with Ca^2+^ bound further support the stabilization of the receptor fold in the presence of an ion in the binding site, as observed in some of the Na^+^-bound simulations. In addition to the lower RMSD values (Figure S5), comparison of the RMSF of the three simulation datasets (Figure S7) further confirms increased stability and reduced flexibility of the receptor with calcium bound, particularly in the TM5-TM6-TM7 region. Analysis of the H-bonds across the five Ca^2+^-bound replicas did not reveal any exclusive interactions that are absent in the trajectories with sodium. Furthermore, in contrast to the simulations with Na^+^, Y291^7.53^ is able to form a hydrogen bond with D69^2.50^ in replicas 2 and 4, but only for about 25% of the trajectory. In replicas 1 and 5, Y291^7.53^ establishes stable hydrophobic interactions with V44^1.53^, L62^2.43^ and I298^8.50^. Generally, in the simulations with Ca^2+^, we observed that the side chain of Y291^7.53^ preferentially orients towards TM1 and TM2. Moreover, the improved electrostatic complementarity between the +2 charge of calcium and the -2 charge of the ion binding site appears to preclude lipid snorkeling, as we did not observe any POPC head groups reaching the ion binding site over the total accumulated 25 *µ*s time. In particular, we noticed a tight coordination of the acidic residues D69^2.50^ and E110^3.39^ with the Ca^2+^ ion (Figure 3c). In this context, visual inspection during the simulation reveals a reduced (almost negligible) amount of water passing between the two sides of the membrane through the receptor (see Table S2).

**Figure 3:**
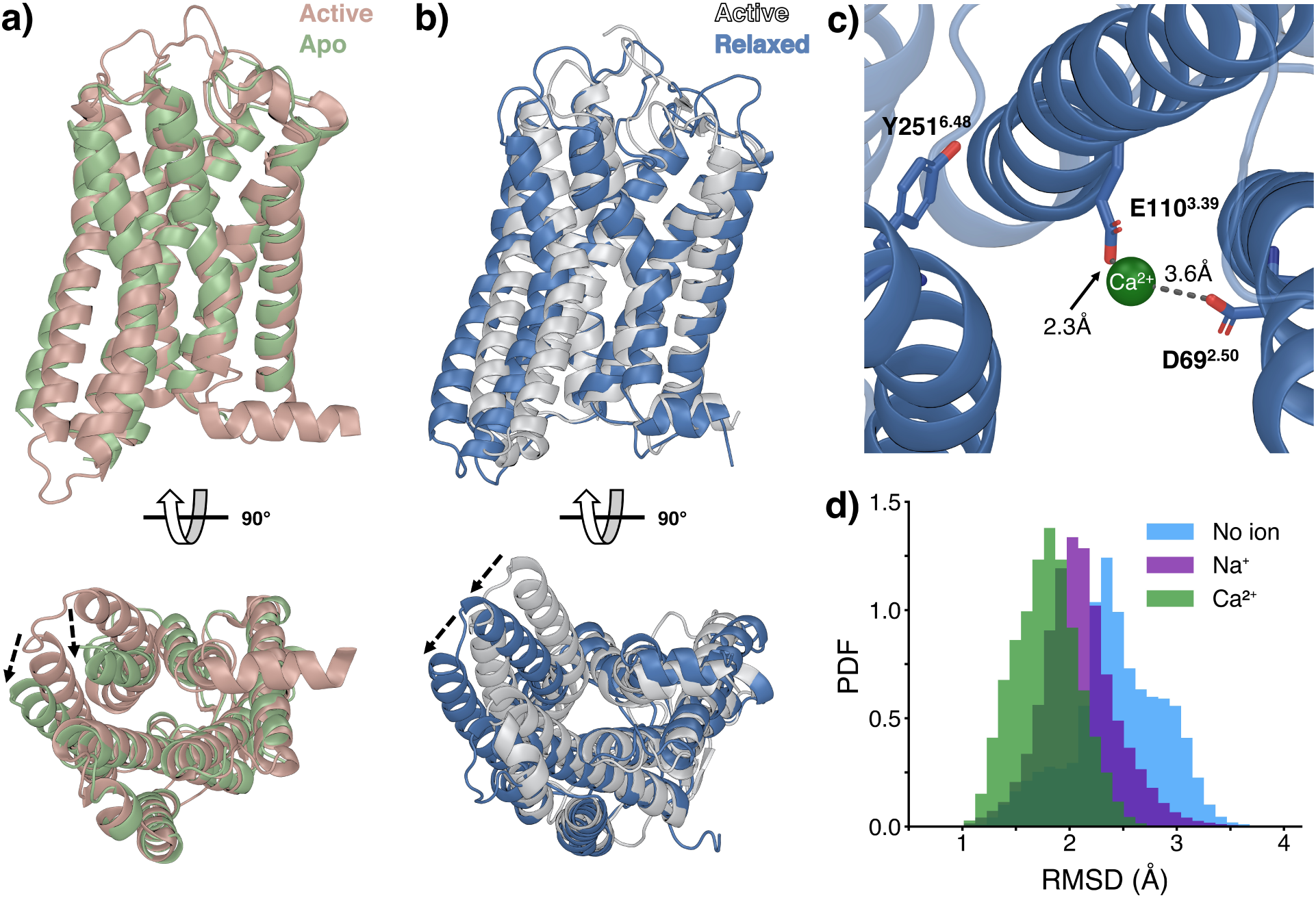
Results of the *in silico* Ca^2+^-bound OR51E2 simulations compared to the available experimental structures. (a) Superposition of the active (orange) and apo (green) structures of OR52_cs_ solved by Choi *et al.*^14^ (b) Comparison of the OR51E2 experimental active structure^7^ (white) and the cluster centroid obtained from the five trajectories with Ca^2+^ in the ion binding site (blue). (c) Detail of the ion binding pocket, highlighting the interactions between Ca^2+^ and the two acidic residues, D69^2.50^ and E110^3.39^. Hydrogen atoms have been omitted for clarity, and the distances refer to the frame shown. (d) Histogram of the RMSD of all the simulations performed in this work (without ions, with Na^+^, and with Ca^2+^ in the ion binding site) with respect to TM3-TM5-TM6 C*_↵_* of the apo form of OR52_cs_.

The main global dynamic effect observed during the Ca^2+^-bound simulations is characterized by the rotational movement of TM5-TM6 around TM3 acting as a pivot (Figure 3b). Such conformational change happens early for all the replicas, during the first 0.3 *÷* 0.7 *µ*s, and appears in agreement with the transition observed for the only experimentally determined active-inactive pair of an OR to date (Figure 3a). Specifically, we performed the calculation of RMSD for all the 75 *µ*s of simulations, taking as reference the C*_↵_* atoms of TM3, TM5, and TM6 of the OR52_cs_ structure, as labeled with the structure alignment tool in the GPCRdb.^24^ We can observe that the RMSD distributions (Figure 3d) are closer to the OR52_cs_ inactive state in the presence of higher-charge ions in the binding site (no ion *>* Na^+^ *>* Ca^2+^). Thus, we can consider the simulations presented in this work as the first observation of calcium-driven *in silico* inactivation of a human OR. Moreover, the TM5-TM6 rotation observed during our simulations for OR51E2 and by comparison of static structures of OR52_cs_ could be considered as the hallmark of inactivation for the OR family.

## Simulations with protonated acidic residues at positions 2.50 and 3.39

To discriminate if receptor inactivation was due simply to neutralization of the doubly charged ion binding pocket by Ca^2+^, we repeated our calculations with a different, neutral protonation state of the two acidic residues D69^2.50^ and E110^3.39^. Operatively, we equilibrated the system with this new protonation state and ran six 1 *µ*s-long independent replicas (that is, longer than the observed timescale for inactivation in the charged system). From the analysis of the trajectories of the neutral system we can observe three main effects. (i) The receptor undergoes a partial active-to-intermediate transition, with a rotation of the TM5-TM6 block, but with a smaller angle with respect to the Ca^2+^-bound charged system (see Figure 4 and Figure S8). (ii) No positively charged ions enter the ion binding site, and (iii) we do not observe any lipid snorkeling by the POPC molecules of the membrane; these last two observations were not unexpected, given the absence of charge imbalance in the ion binding pocket upon protonation of the acidic residues. The absence of ion binding is in line with a previous work on non-olfactory class A GPCRs, where the protonation of the single one acidic residue in the ion pocket facilitates the unbinding of Na^+^.^19^ Taken together, these data indicate that it is not only charge neutralization of the two acidic residues in the ion binding site that drives the transition to the inactive state, but also requires the presence of a divalent ion. This observation is within the limitations of the *µ*s timescale of our simulations and the force field representation of divalent ions and fixed protonation states.

**Figure 4:**
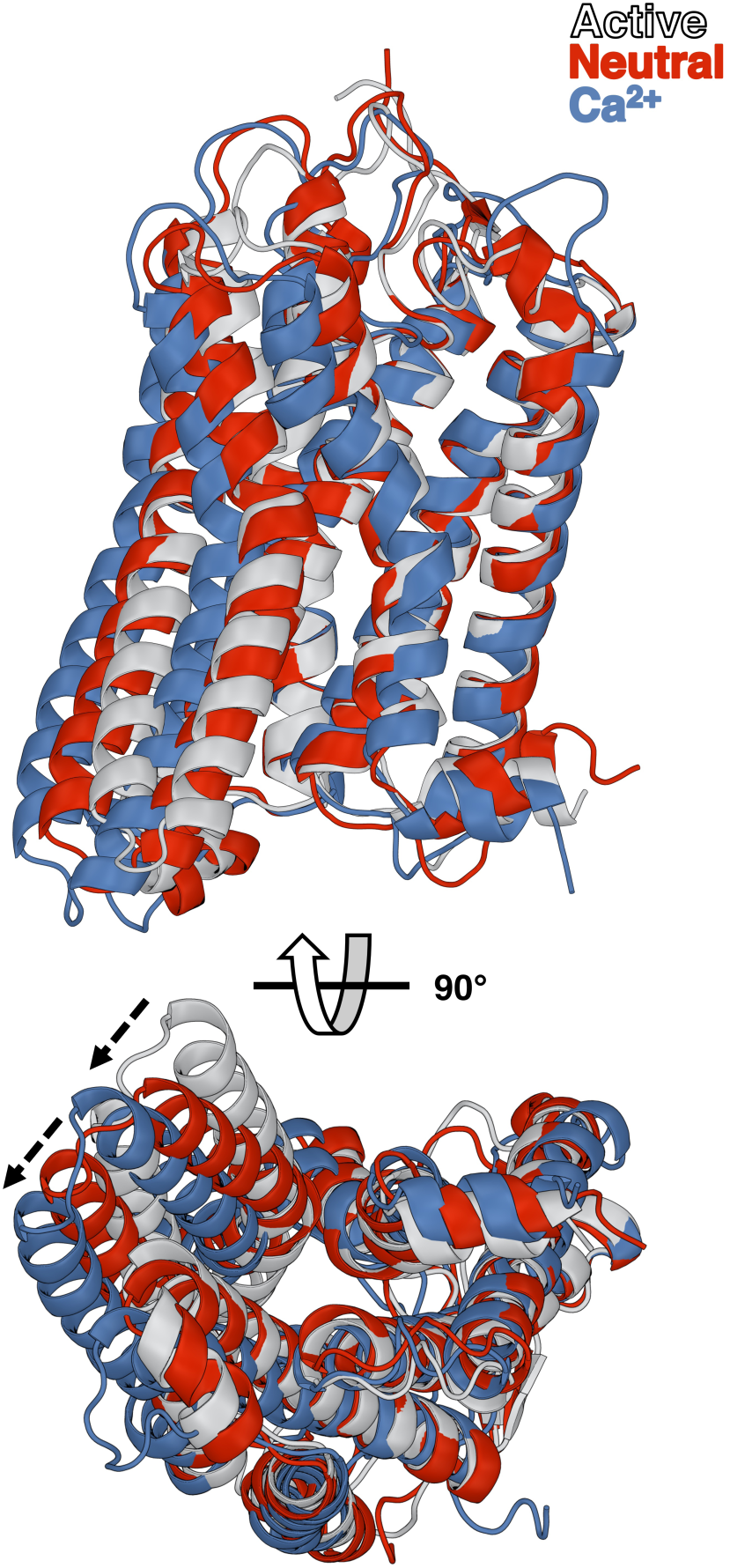
Results of the simulations with protonated residues at positions 2.50 and 3.39. Superposition of the cryoEM structure in its active state (white), the cluster centroid obtained from the five trajectories with Ca^2+^ bound (blue), and the cluster centroid obtained from the six trajectories with neutral D^2.50^ and E^3.39^ (red). The progression of TM5-TM6 rotation in the three structures is indicated with dashed arrows.

## Discussion

The conservation of two acidic residues in the ion binding site of ~ 98% of human ORs,^13^ at positions 2.50 and 3.39 (Figure 5a), raises the question of whether not only monovalent, but also ions divalent, can occupy this site and modulate receptor function. The simulations carried out in this work show that indeed calcium is more efficient than sodium at stabilizing the inactive state of a prototypical OR, OR51E2. Neutralization of the ion binding site by protonation of the two acidic residues is not sufficient to drive receptor inactivation, further supporting the strict requirement for a divalent ion bound in this pocket to stabilize the inactive receptor state.

**Figure 5:**
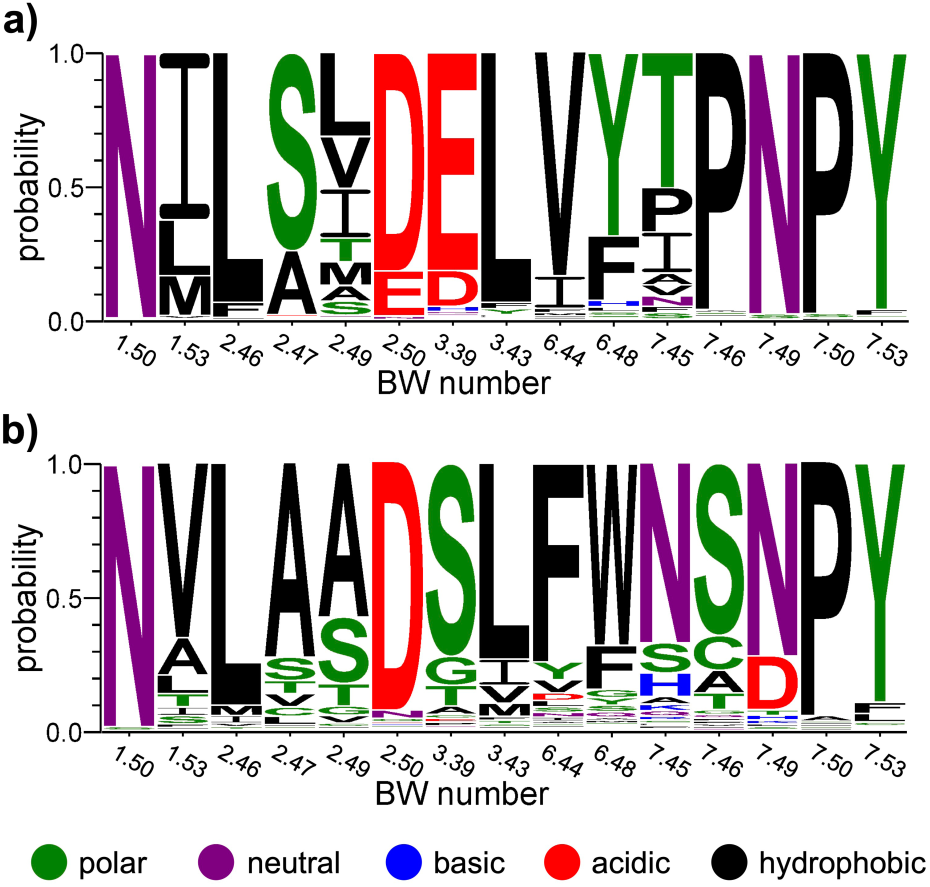
Sequence logos depicting conservation of residues lining the ion binding site in class A GPCRs. (a) Human olfactory receptors, based on the multiple sequence alignment of 412 sequences in Fierro *et al.*^36^ (b) Human non-olfactory class A GPCRs, using the 286 sequences available in the GPCRdb sequence alignment tool^37^ as of January 2024. Sequence logos were generated with WebLogo version 3.7.12^38^ and amino acids are colored according to their chemical properties.

The possibility of divalent ions binding to class A GPCRs containing two acidic residues in the ion binding site was already highlighted in the review by Katritch *et al.* as an “outstanding question”. ^23^ Besides the conserved D^2.50^, previous sequence analyses^23,25^ have shown that a subset of non-olfactory class A GPCRs bear a second acidic residue in the ion binding site. Here we extended such analyses and estimated the number to be ~ 21%. The second acidic residue is typically located at position 7.49 (~ 18%),but we also observed a nonnegligible number of receptors where it is located at positions 6.44 (~ 2%) and 3.39 (~ 1%) (see Figure 5b and Table S3). However, as of January 2024 only the experimental structures of melanocortin receptors^26–32^ have shown Ca^2+^ bound to two aspartates at positions 2.50 and 7.49, acting as a cofactor for agonist binding. Instead, experimental structures of proteinase-activated receptors^33,34^ and cysteinyl leukotriene receptors, ^35^ also bearing two aspartates at positions 2.50 and 7.49, have been solved with Na^+^ and an antagonist bound.

We thus believe that the investigation of calcium ions effect is an extremely promising venue in the context of GPCRs,^39^ in particular for ORs, which also needs an experimental counterpart. Despite the great advancements in the experimental determination of GPCR structures, the state of the art in cryo-EM is still unable to discriminate between water and ions.^40^ A possible help may come from the use of buffers with high(er) calcium concentrations, focusing our attention on possible ion-protein interactions. Moreover, while most non-olfactory class A GPCRs with a second acidic residue (see Table S3) bear an aspartate at either position 7.49 (~ 86%) or 6.44 (~ 8%), the majority of ORs (~ 90%) have longer side chain acidic residue (glutamate) and at different position (3.39).^13^ This is expected to shift the ion location within the binding site, thus resulting in di↵erent e↵ects of ion binding on receptor function. Interestingly, two chemokine receptors (CCR1 and CCR3) and one peptide receptor (QRFP) also display a glutamate at position 3.39, suggesting the possibility that calcium might also stabilize the inactive state of these receptors, similarly to what we proposed here for ORs.

In summary, in this work we suggest a molecular mechanism for ORs inactivation, which relies on the presence of calcium in the ion binding site. In this regard, calcium-mediated OR inactivation could constitute a negative feedback mechanism to stop the olfactory signal triggered by odorant binding and subsequent calcium entry through cyclic nucleotide gated channels. The work shown here suffers the technical limitations of classical modeling approaches in the presence of divalent ions and fixed protonation states. We envisage the possibility to use more advanced techniques such as QM/MM, which proved their efficacy in explaining classical model shortcomings in polarization effects in GPCRs,^41^ exploiting the next generation exascale HPC machines. We hope that this computationally-driven hypotheses could be validated by means of experimental techniques, highlighting the predictive power of molecular modeling and simulations to suggest biologically relevant structure-function relationships. Such experimental testing could include *in vitro* assays with varying concentrations of calcium in the extracellular medium or structural determination efforts with higher calcium (or divalent metal analogs) concentrations in the buffers used for X-ray, cryo-EM or native mass spectrometry.^42^

## Methods

### System preparation

The initial configuration of OR51E2 in its active state was obtained from the Protein Data Bank (PDB code: 8F76^7^). This structure was preprocessed using the Protein Preparation Wizard implemented in Schrödinger Maestro 2023-3,^43^ which automatically assigns the amino acid protonation states. In particular, both D69^2.50^ and E110^3.39^ were predicted to be negatively charged at pH 7.4 and in the presence of a positively charged ion (either Na^+^ or Ca^2+^) in the ion binding site. Instead, in the absence of ions both D69^2.50^ and E110^3.39^ were predicted to be protonated. We refer to these two alternative protonation states as “charged” and “neutral” forms of the receptor, respectively. The complete system was then assembled using the CHARMM-GUI^20,21^ webserver. First, a disulfide bond was established between C96^3.25^ and C178^45.50^; then, a cubic box of dimensions 100 x 100 x 120Å was defined with the receptor embedded in a mixed lipid bilayer composed of POPC and cholesterol (3:1 ratio). The receptor-membrane system was then solvated in water with a NaCl concentration of 0.15 M. In the case of the calcium-bound simulations, we added a single Ca^2+^ ion and neutralized its charge by adding two additional Cl^-^ ions in the solution. The protein, lipids, and ions were parameterized using the CHARMM36m force field,^44^ while water was described using the TIP3P^45^ model.

### Molecular dynamics simulations

In this work, the equilibration phase of the simulations was conducted following a protocol presented in our previous work;^13^ further details are provided in the Supporting Information. For the subsequent production phase, we performed unrestrained molecular dynamics (MD), with a 2 fs time step. Simulations were 5 *µ*s long for the three charged systems and 1 *µ*s for the neutral apo form. To maintain constant temperature and pressure conditions at 310 K and 1 bar, respectively, we employed the velocity rescaling thermostat^46^ alongside the semi-isotropic cell rescaling barostat. ^47^ A total of 21 independent runs were executed, comprising (i) five replicates for each of the three charged models, i.e. without any ions in the ion binding pocket, with Na^+^ bound, and with Ca^2+^ bound and (ii) six replicates of the neutral model without ions in the ion binding pocket. Each of these runs was initiated with distinct initial velocities. The simulations were all carried out using GROMACS,^48^ version 2021.2.

## Data Availability Statement

Data needed to reproduce the results shown in this paper (structures, topology, GROMACS input files, etc.) and the resulting trajectories are available at Zenodo (https://zenodo.org/ doi/10.5281/zenodo.10589509). All the trajectories computed here for OR51E2 with charged D^2.50^ and E^3.39^ have been also uploaded on the GPCRmd,^49^ with the accession codes 1976 (no ions), 1977 (Na^+^-bound), and 1978 (Ca^2+^-bound).

## Acknowledgement

We thank Prof. Carlo Camilloni and Dr. Antonella Paladino for fruitful discussion and Prof. Hee-Jung Choi for providing us, before its official deposit in the Protein Data Bank, the apo structure of OR52_cs_. We also thank the anonymous reviewers for helping us in strengthening and improving the work presented here. We acknowledge EuroHPC Joint Undertaking for awarding access to the Karolina CPU cluster at IT4Innovations, Czech Republic (EHPC-REG-2022R03-049). We also acknowledge the CINECA award under the ISCRA initiative, for the availability of high-performance computing resources and support. This research was supported by Università degli Studi di Milano (Piano di supporto di ateneo Linea 2 2023-DBS Capelli) to R.C. and in part by the DFG Research Unit FOR2518 “Functional Dynamics of Ion Channels and Transporters – DynIon”, project P6, to M.A.-P.

## Supporting Information Available

The supporting information file contains the extended details of the MD simulations and tables with the Ballesteros-Weinstein numbering of OR51E2, with the main events observed in every MD replica of the charged systems and with the list of non-olfactory class A GPCRs with two acidic residues in the ion binding site, as well as figures showing the RMSD of all the replicas of the charged systems with respect to the experimental structure of OR51E2 in the active state, the minimum distance analysis between POPC lipid molecules and the ion binding site, and the RMSF for all the simulations performed for the three charged systems, and RMSD histograms with respect to OR52_cs_ for all systems, either charged or neutral.

