## Supporting Information file for "Calcium-driven In Silico Inactivation of a Human Olfactory Receptor"

#### Preparation of the Structural Models

In the initial phase, the cryoEM structure of OR51E2 in its active state (PDB code: 8F76[1]) was prepared using the Protein Preparation Wizard in Schrödinger Maestro version 2023-3.[2] This process involved automated assignment of protonation states for amino acids based on their microenvironment. D69$^{2.50}$ and E110$^{3.39}$ were automatically identified as negatively charged in the presence of a $Na^+$ or $Ca^{2+}$ ion. In absence of any ions, both D69$^{2.50}$ and E110$^{3.39}$ were instead predicted to be protonated. This dependency of the protonation state of the ion binding site on the presence of ions has also been reported for non-olfactory class A receptors with a single acidic residue at position 2.50.[3] The thus-prepared models (neutral apo receptor and charged receptor in either apo, $Na^{+}$-bound and $Ca^{2+}$-bound forms) were then subjected to further refinement using CHARMM-GUI online platform[4,5] (https://charmm-gui.org/). This included establishing a disulfide bond between $C96^{3.25}$ and $C178^{45.50}$ and setting up a cubic simulation box of dimensions (100 Å) × (100 Å) × (120 Å). Within this box, the receptor was located at the center, embedded in a membrane comprising a 3:1 POPC:cholesterol mix. The system, including the lipid bilayer and receptor, was immersed in water with 150 mM NaCl, reflecting standard experimental conditions for GPCRs. In the replicas with the calcium ion bound, we also added two chloride ions to neutralize the extra 2+ charge. The CHARMM36m force field[6] was used to parameterize proteins, lipids, and ions, while water molecules were modeled using the TIP3P[7] model. When necessary, we manually positioned the sodium or calcium ion in the ion binding pocket of the olfactory receptor.

### Molecular Dynamics Simulation Protocol

The molecular dynamics simulation protocol adopted was the same as the one presented in our previous work on inactive models of olfactory receptors.[8] In practice, it is an adaptation and extension of the standard procedure recommended by CHARMM-GUI for transmembrane proteins. It includes nine distinct steps (according to the CHARMM-GUI numbering system, from 0 to 6, plus two additional phases, 7 and 8):

- step 0: 5,000 steps of steepest descent minimization, constraining the protein backbone ($k = 4,000$ kJ/mol/nm$^2$) and side chains ($k = 2,000$ kJ/mol/nm$^2$), lipid phosphate groups ($k = 1,000$ kJ/mol/nm$^2$), and dihedrals ($k = 1,000$ kJ/mol/rad$^2$).
- Step 1: 125 ps of MD with a time step of 1 fs, restraining the protein backbone ($k = 4,000$ kJ/mol/nm$^2$) and side chains ($k = 2,000$ kJ/mol/nm$^2$), phosphate groups for POPC and hydroxyl group for cholesterol ($k = 1,000$ kJ/mol/nm$^2$), and dihedrals ($k = 1,000$ kJ/mol/rad$^2$).

- Step 2: 125 ps of MD with a time step of 1 fs, restraining the protein backbone ($k = 2,000$ kJ/mol/nm$^2$) and side chains ($k = 1,000$ kJ/mol/nm$^2$), phosphate groups for POPC and hydroxyl group for cholesterol ($k = 400$ kJ/mol/nm$^2$), and dihedrals ($k = 400$ kJ/mol/rad$^2$).

- Step 3: 125 ps of MD with a time step of 1 fs, restraining the protein backbone ($k = 1,000$ kJ/mol/nm$^2$) and side chains ($k = 500$ kJ/mol/nm$^2$), phosphate groups for POPC and hydroxyl group for cholesterol ($k = 400$ kJ/mol/nm$^2$) and dihedrals ($k = 200$ kJ/mol/rad$^2$).

- Step 4: 500 ps MD with a time step of 2 fs, restraining the protein backbone ($k = 500$ kJ/mol/nm$^2$) and side chains ($k = 200$ kJ/mol/nm$^2$), phosphate groups for POPC and hydroxyl group for cholesterol ($k = 200$ kJ/mol/nm$^2$) and dihedrals ($k = 200$ kJ/mol/rad$^2$).

- Step 5: 500 ps of MD with a time step of 2 fs, restraining the protein backbone ($k = 200$ kJ/mol/nm$^2$) and side chains ($k = 50$ kJ/mol/nm$^2$), phosphate groups for POPC and hydroxyl group for cholesterol ($k = 40$ kJ/mol/nm$^2$), and dihedrals ($k = 100$ kJ/mol/rad$^2$).

- Step 6: 100 ns MD with a time step of 2 fs, restraining the protein backbone ($k = 50$ kJ/mol/nm$^2$); this step is 10 times longer than the standard CHARMM-GUI protocol.

- Step 7: 100 ns of MD with a time step of 2 fs, restraining the protein backbone ($k = 5$ kJ/mol/nm$^2$); this is a completely new step that increases the length of the restrained equilibration.

The final production phase (step 8) consisted of MD without any restraint, 5 $\mu$s long for the charged systems and 1 $\mu$s for the neutral apo receptor. All simulations were performed with a 2 fs time step. The cutoff for van der Waals and short-range interactions was set to 10 Å , and long-range electrostatic interactions were calculated using the Ewald smooth particle mesh method.[9] Temperature control was achieved using the velocity rescale thermostat[10] at 310 K, and pressure was maintained at 1 bar using the semi-isotropic cell rescale barostat.[11] All simulations were performed with GROMACS[12] version 2021.2.

In total, we ran fifteen different production simulations: five replicas starting with no ions in the binding pocket, five starting with sodium in the ion binding pocket, and five starting with calcium in the binding pocket. Each simulation with charged D69$^{2.50}$ and E110$^{3.39}$ was 5 $\mu$s-long, while the replicas with protonated acidic residues were 1 $\mu$s-long.

### Hydrogen Bonds Analysis

The 15 trajectories were analyzed using the 'Hydrogen Bonds' plugin from the Visual Molecular Dynamics[13] (VMD) suite, version 1.9.3. The analysis was configured to identify hydrogen bonds involving only polar atoms of the receptor, considering a donor-acceptor distance within 3.5 Å, and a tolerance of 30° deviation from the linear donor-hydrogen-acceptor angle. For comparison among replicas, a custom Python3[14] script was developed aimed at identifying hydrogen bonds in a selected trajectory that were not present in all the other trajectories, given a specified threshold for the hydrogen bond occupancy percentage.

### Cluster Analysis

Cluster analyses were performed on the concatenated trajectories of the five replicas for each simulation condition (no ions, $Na^+$, and $Ca^{2+}$, respectively).

The clustering was performed using the gromos method[15] implemented in the GROMACS cluster tool, setting an RMSD cutoff of 4 Å. The RMSD was calculated on the $C_\alpha$ atoms of the transmembrane helices (TM1 to TM7), as defined in Table S1.

#### RMSD and RMSF analyses

All the RMSD and RMSF analyses shown here are performed on the $C_\alpha$ of the transmembrane helices of the OR51E2 (see Table S1). For the RMSD, we consider the cryoEM structure of the receptor in its active state[1] as the reference conformation. The RMSF calculations (Figure S8) have been performed on the concatenated trajectories for each the three simulation conditions, encompassing 25 $\mu$s of dynamics each.

Table S1: Ballesteros-Weinstein generic numbering for human OR51E2, as listed in the GPCRdb[16] (https://gpcrdb.org/residue/residuetabledisplay, accessed on December 2023).

| **TM1** | | **TM2** | | **TM3** | | **TM4** | | **TM5** | | **TM6** | | **TM7** | |
|---|---|---|---|---|---|---|---|---|---|---|---|---|---|
| 1x32 | H23 | 2x37 | A56 | 3x21 | S92 | 4x38 | N136 | 5x32 | T191 | 6x26 | S229 | 7x29 | H268 |
| 1x33 | F24 | 2x38 | P57 | 3x22 | F93 | 4x39 | N137 | 5x33 | L192 | 6x27 | K230 | 7x30 | P269 |
| 1x34 | W25 | 2x39 | M58 | 3x23 | E94 | 4x40 | T138 | 5x34 | P193 | 6x28 | S231 | 7x31 | I270 |
| 1x35 | V26 | 2x40 | Y59 | 3x24 | A95 | 4x41 | V139 | 5x35 | N194 | 6x29 | E232 | 7x32 | V271 |
| 1x36 | G27 | 2x41 | L60 | 3x25 | C96 | 4x42 | T140 | 5x36 | V195 | 6x30 | R233 | 7x33 | R272 |
| 1x37 | F28 | 2x42 | F61 | 3x26 | L97 | 4x43 | A141 | 5x37 | V196 | 6x31 | A234 | 7x34 | V273 |
| 1x38 | P29 | 2x43 | L62 | 3x27 | T98 | 4x44 | Q142 | 5x38 | Y197 | 6x32 | K235 | 7x35 | V274 |
| 1x39 | L30 | 2x44 | C63 | 3x28 | Q99 | 4x45 | I143 | 5x39 | G198 | 6x33 | A236 | 7x36 | M275 |
| 1x40 | L31 | 2x45 | M64 | 3x29 | M100 | 4x46 | G144 | 5x40 | L199 | 6x34 | F237 | 7x37 | G276 |
| 1x41 | S32 | 2x46 | L65 | 3x30 | F101 | 4x47 | I145 | 5x41 | T200 | 6x35 | G238 | 7x38 | D277 |
| 1x42 | M33 | 2x47 | A66 | 3x31 | F102 | 4x48 | V146 | 5x42 | A201 | 6x36 | T239 | 7x39 | I278 |
| 1x43 | Y34 | 2x48 | A67 | 3x32 | I103 | 4x49 | A147 | 5x43 | I202 | 6x37 | C240 | 7x40 | Y279 |
| 1x44 | V35 | 2x49 | I68 | 3x33 | H104 | 4x50 | V148 | 5x44 | L203 | 6x38 | V241 | 7x41 | L280 |
| 1x45 | V36 | 2x50 | D69 | 3x34 | A105 | 4x51 | V149 | 5x45 | L204 | 6x39 | S242 | 7x42 | L281 |
| 1x46 | A37 | 2x51 | L70 | 3x35 | L106 | 4x52 | R150 | 5x46 | V205 | 6x40 | H243 | 7x43 | L282 |
| 1x47 | M38 | 2x52 | A71 | 3x36 | S107 | 4x53 | G151 | 5x47 | M206 | 6x41 | I244 | 7x45 | P283 |
| 1x48 | F39 | 2x53 | L72 | 3x37 | A108 | 4x54 | S152 | 5x48 | G207 | 6x42 | G245 | 7x46 | P284 |
| 1x49 | G40 | 2x54 | S73 | 3x38 | I109 | 4x55 | L153 | 5x49 | V208 | 6x43 | V246 | 7x47 | V285 |
| 1x50 | N41 | 2x55 | T74 | 3x39 | E110 | 4x56 | F154 | 5x50 | D209 | 6x44 | V247 | 7x48 | I286 |
| 1x51 | C42 | 2x551 | S75 | 3x40 | S111 | 4x57 | F155 | 5x51 | V210 | 6x45 | L248 | 7x49 | N287 |
| 1x52 | I43 | 2x56 | T76 | 3x41 | T112 | 4x58 | F156 | 5x52 | M211 | 6x46 | A249 | 7x50 | P288 |
| 1x53 | V44 | 2x57 | M77 | 3x42 | I113 | 4x59 | P157 | 5x53 | F212 | 6x47 | F250 | 7x51 | I289 |
| 1x54 | V45 | 2x58 | P78 | 3x43 | L114 | 4x60 | L158 | 5x54 | I213 | 6x48 | Y251 | 7x52 | I290 |
| 1x55 | F46 | 2x59 | K79 | 3x44 | L115 | 4x61 | P159 | 5x55 | S214 | 6x49 | V252 | 7x53 | Y291 |
| 1x56 | I47 | 2x60 | I80 | 3x45 | A116 | 4x62 | L160 | 5x56 | L215 | 6x50 | P253 | 7x54 | G292 |
| 1x57 | V48 | 2x61 | L81 | 3x46 | M117 | 4x63 | L161 | 5x57 | S216 | 6x51 | L254 | 7x55 | A293 |
| 1x58 | R49 | 2x62 | A82 | 3x47 | A118 | 4x64 | I162 | 5x58 | Y217 | 6x52 | I255 | 7x56 | K294 |
| 1x59 | T50 | 2x63 | L83 | 3x48 | F119 | 4x65 | K163 | 5x59 | F218 | 6x53 | G256 | | |
| 1x60 | E51 | 2x64 | F84 | 3x49 | D120 | 4x66 | R164 | 5x60 | L219 | 6x54 | L257 | | |
| | | 2x65 | W85 | 3x50 | R121 | 4x67 | L165 | 5x61 | I220 | 6x55 | S258 | | |
| | | 2x66 | F86 | 3x51 | Y122 | | | 5x62 | I221 | 6x56 | V259 | | |
| | | 2x67 | D87 | 3x52 | V123 | | | 5x63 | R222 | 6x57 | V260 | | |
| | | | | 3x53 | A124 | | | 5x64 | T223 | 6x58 | H261 | | |
| | | | | 3x54 | I125 | | | 5x65 | V224 | 6x59 | R262 | | |
| | | | | 3x55 | C126 | | | 5x66 | L225 | 6x60 | F263 | | |
| | | | | 3x56 | H127 | | | 5x67 | Q226 | 6x61 | G264 | | |
| | | | | | | | | 5x68 | L227 | | | | |

| **ICL1** | | | | **ICL2** | | **ECL2** | | | | | | **H8** | |
|---|---|---|---|---|---|---|---|---|---|---|---|---|---|
| 12x48 | R52 | | | 34x50 | P128 | 45x50 | C178 | | | | | 8x47 | T295 |
| 12x49 | S53 | | | 34x51 | L129 | 45x51 | V179 | | | | | 8x48 | K296 |
| 12x50 | L54 | | | 34x52 | R130 | 45x52 | H180 | | | | | 8x49 | Q297 |
| 12x51 | H55 | | | 34x53 | H131 | | | | | | | 8x50 | I298 |
| | | | | 34x54 | A132 | | | | | | | 8x51 | R299 |
| | | | | 34x55 | A133 | | | | | | | 8x52 | T300 |
| | | | | 34x56 | V134 | | | | | | | 8x53 | R301 |
| | | | | 34x57 | L135 | | | | | | | 8x54 | V302 |
| | | | | | | | | | | | | 8x55 | L303 |
| | | | | | | | | | | | | 8x56 | A304 |
| | | | | | | | | | | | | 8x57 | M305 |
| | | | | | | | | | | | | 8x58 | F306 |
| | | | | | | | | | | | | 8x59 | K307 |
| | | | | | | | | | | | | 8x60 | I308 |

Table S2: Relevant events observed in each replica.

| Initial condition | Replica ID | Ion un/binding | POPC snorkeling | Role of $Y^{6.48}$ | Role of $Y^{7.53}$ | Water passing |
|---|---|---|---|---|---|---|
| No ion | 1 | Binding at 0.44 $\mu$s | 1 POPC from 0.44 $\mu$s | H-bond with $S^{3.40}$ | Interacts with $V^{1.53}$, $L^{2.63}$, $I^{8.50}$ from 0.17 to 1 $\mu$s and from 2 to 2.2 $\mu$s | Free passage |
| | 2 | No | 2 POPC: 1$^{st}$ from 0.26 $\mu$s and 2$^{nd}$ from 1.49 $\mu$s | Interacts with POPC | Interacts with $V^{1.53}$, $L^{2.63}$, $I^{8.50}$ from 1.34 to 2.17 $\mu$s; then with POPC | Free passage |
| | 3 | No | 1 POPC from 0.09 $\mu$s | Interacts with POPC | Interacts with POPC | Free passage |
| | 4 | Binding at 4.66 $\mu$s | 1 POPC from 0.03 $\mu$s to 4.64 $\mu$s | H-bond with $E^{3.39}$ from 2.35 $\mu$s and $S^{3.40}$ from 4.49 $\mu$s | Interacts with POPC | Free passage |
| | 5 | No | 1 POPC from 0.03 $\mu$s | Interacts with $E^{3.39}$, and with POPC from 0.03 $\mu$s | Interacts with POPC | Free passage |
| $Na^{+}$ | 1 | No | No | H-bond with $S^{3.40}$ until 2.10 $\mu$s and with $E^{3.39}$ from 2.10 $\mu$s | Interacts with $M^{3.46}$, $I^{8.50}$ and from 1.7 $\mu$s with $D^{2.50}$ | Free passage |
| | 2 | No | 1 POPC from 4.14 $\mu$s | H-bond with $S^{3.40}$ | Interacts with $D^{2.50}$ | Few molecules |
| | 3 | No | No | H-bond with $S^{3.40}$ | Interacts with $D^{2.50}$ | Free passage |
| | 4 | No | 1 POPC from 0.15 $\mu$s to 2.15 $\mu$s | H-bond with $S^{3.40}$ | Points towards TM3-TM5 | Free passage |
| | 5 | Unbinding at 4.2 $\mu$s, and binding at 4.95 $\mu$s | 1 POPC from 0.76 $\mu$s to 3.15 $\mu$s 2 POPC: 1$^{st}$ from 3.83 $\mu$s and 2$^{nd}$ from 4.23 $\mu$s | H-bond with $S^{3.40}$ until 1.85 $\mu$s | Interacts with POPC | Free passage |
| $Ca^{2+}$ | 1 | No | No | H-bond with $S^{3.40}$ | Interacts with $V^{1.53}$, $L^{2.63}$, $I^{8.50}$ from 1.25 $\mu$s | Few molecules |
| | 2 | No | No | H-bond with $S^{3.40}$ | Interacts with $D^{2.50}$ from 0.2 to 2.38 $\mu$s | Almost water-free |
| | 3 | No | No | H-bond with $S^{3.40}$ | Interacts with $D^{2.50}$ from 2.9 to 3.5 $\mu$s | Almost water-free |
| | 4 | No | No | H-bond with $S^{3.40}$ | Interacts with $D^{2.50}$ multiple times | Few molecules |
| | 5 | No | No | H-bond with $S^{3.40}$ | Interacts with $V^{1.53}$, $L^{2.63}$, $I^{8.50}$ from 0.37 $\mu$s | Almost water-free |

Table S3: Non-olfactory class A GPCRs with a second acidic residue in the ion binding site, besides the first acidic residue at position 2.50. Out of the 286 human sequences available in the sequence alignment tool of GPCRdb[17] as of January 2024, 60 bear a second acidic residue in the ion binding site, specifically, $E^{3.39}$ in three receptors, $D^{6.44}$ in five and $D^{7.49}$ in 55.

| Uniprot ID | First | Second | Uniprot ID | First | Second |
|---|---|---|---|---|---|
| acthr_human | $D^{2.50}$ | $D^{7.49}$ | mshr_human | $D^{2.50}$ | $D^{7.49}$ |
| ccr1_human | $D^{2.50}$ | $E^{3.39}$ | ogr1_human | $D^{2.50}$ | $D^{7.49}$ |
| ccr3_human | $D^{2.50}$ | $E^{3.39}$ | oxer1_human | $D^{2.50}$ | $D^{7.49}$ |
| cltr1_human | $D^{2.50}$ | $D^{7.49}$ | p2ry1_human | $D^{2.50}$ | $D^{7.49}$ |
| ffar2_human | $D^{2.50}$ | $D^{7.49}$ | p2ry2_human | $D^{2.50}$ | $D^{7.49}$ |
| ffar3_human | $D^{2.50}$ | $D^{7.49}$ | p2ry4_human | $D^{2.50}$ | $D^{7.49}$ |
| fshr_human | $D^{2.50}$ | $D^{6.44}$ | p2ry6_human | $D^{2.50}$ | $D^{7.49}$ |
| gp132_human | $E^{2.50}$ | $D^{7.49}$ | p2ry8_human | $D^{2.50}$ | $D^{7.49}$ |
| gp171_human | $D^{2.50}$ | $D^{7.49}$ | p2y10_human | $D^{2.50}$ | $D^{7.49}$ |
| gp174_human | $D^{2.50}$ | $D^{7.49}$ | p2y12_human | $D^{2.50}$ | $D^{7.49}$ |
| gp183_human | $D^{2.50}$ | $D^{7.49}$ | p2y13_human | $D^{2.50}$ | $D^{7.49}$ |
| gpr17_human | $D^{2.50}$ | $D^{7.49}$ | p2y14_human | $D^{2.50}$ | $D^{7.49}$ |
| gpr18_human | $D^{2.50}$ | $D^{7.49}$ | par1_human | $D^{2.50}$ | $D^{7.49}$ |
| gpr20_human | $D^{2.50}$ | $D^{7.49}$ | par2_human | $D^{2.50}$ | $D^{7.49}$ |
| gpr34_human | $D^{2.50}$ | $D^{7.49}$ | par3_human | $D^{2.50}$ | $D^{7.49}$ |
| gpr35_human | $D^{2.50}$ | $D^{7.49}$ | par4_human | $D^{2.50}$ | $D^{7.49}$ |
| gpr42_human | $D^{2.50}$ | $D^{7.49}$ | pd2r_human | $D^{2.50}$ | $D^{7.49}$ |
| gpr4_human | $D^{2.50}$ | $D^{7.49}$ | pe2r1_human | $D^{2.50}$ | $D^{7.49}$ |
| gpr55_human | $D^{2.50}$ | $D^{7.49}$ | pe2r2_human | $D^{2.50}$ | $D^{7.49}$ |
| gpr87_human | $D^{2.50}$ | $D^{7.49}$ | pe2r3_human | $D^{2.50}$ | $D^{7.49}$ |
| hcar1_human | $D^{2.50}$ | $D^{7.49}$ | pe2r4_human | $D^{2.50}$ | $D^{7.49}$ |
| hcar2_human | $D^{2.50}$ | $D^{7.49}$ | pf2r_human | $D^{2.50}$ | $D^{7.49}$ |
| hcar3_human | $D^{2.50}$ | $D^{7.49}$ | pi2r_human | $D^{2.50}$ | $D^{7.49}$ |
| lpar4_human | $D^{2.50}$ | $D^{7.49}$ | psyr_human | $D^{2.50}$ | $D^{7.49}$ |
| lpar5_human | $D^{2.50}$ | $D^{7.49}$ | ptafr_human | $D^{2.50}$ | $D^{7.49}$ |
| lpar6_human | $D^{2.50}$ | $D^{7.49}$ | qrfpr_human | $D^{2.50}$ | $E^{3.39}$ |
| lshr_human | $D^{2.50}$ | $D^{6.44}$ | rxfp1_human | $D^{2.50}$ | $D^{6.44}$ |
| mc3r_human | $D^{2.50}$ | $D^{7.49}$ | rxfp2_human | $D^{2.50}$ | $D^{6.44}$ |
| mc4r_human | $D^{2.50}$ | $D^{7.49}$ | ta2r_human | $D^{2.50}$ | $D^{7.49}$ |
| mc5r_human | $D^{2.50}$ | $D^{7.49}$ | tshr_human | $D^{2.50}$ | $D^{6.44}$ |

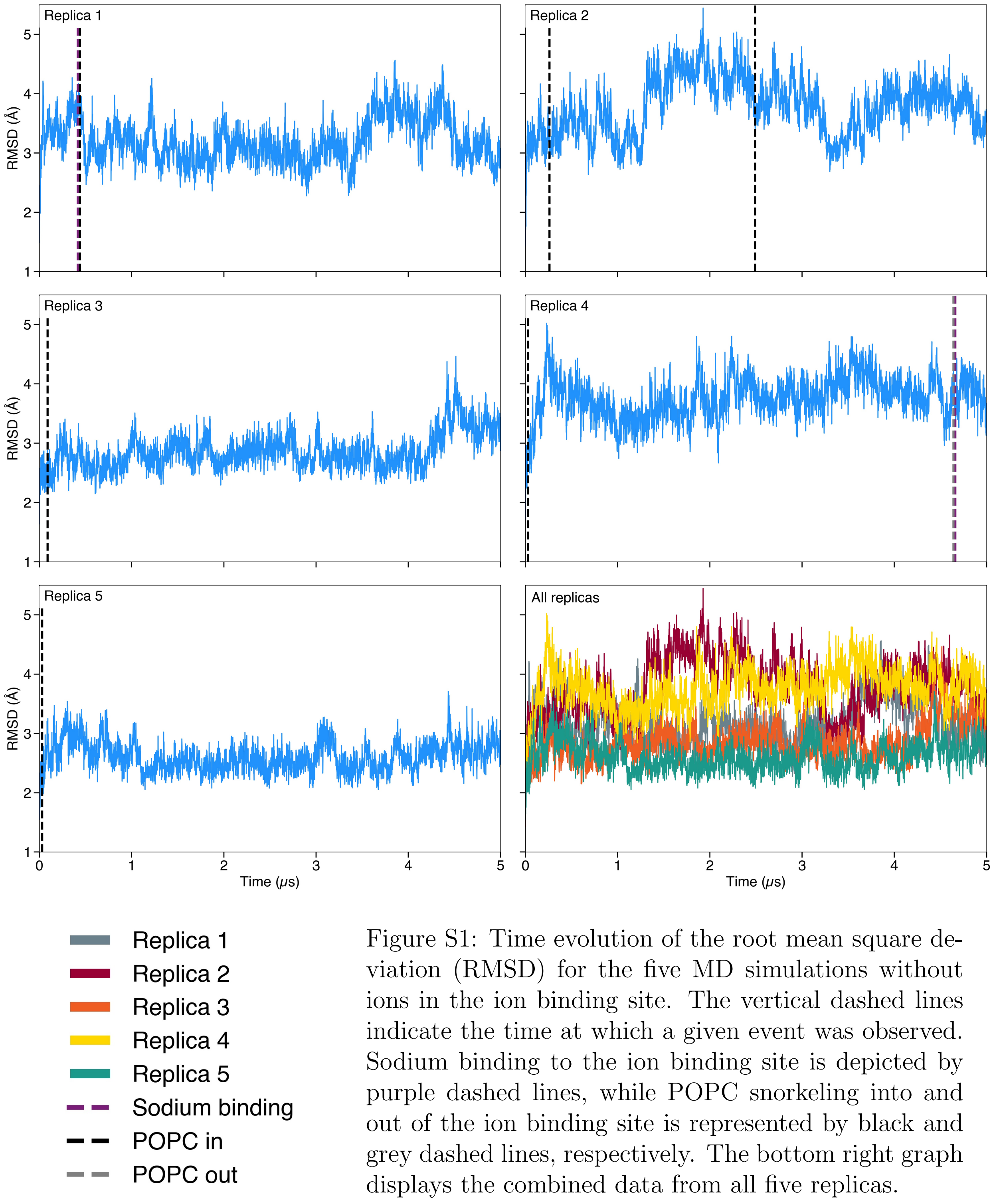


Figure S1: Time evolution of the root mean square deviation (RMSD) for the five MD simulations without ions in the ion binding site. The vertical dashed lines indicate the time at which a given event was observed. Sodium binding to the ion binding site is depicted by purple dashed lines, while POPC snorkeling into and out of the ion binding site is represented by black and grey dashed lines, respectively. The bottom right graph displays the combined data from all five replicas.

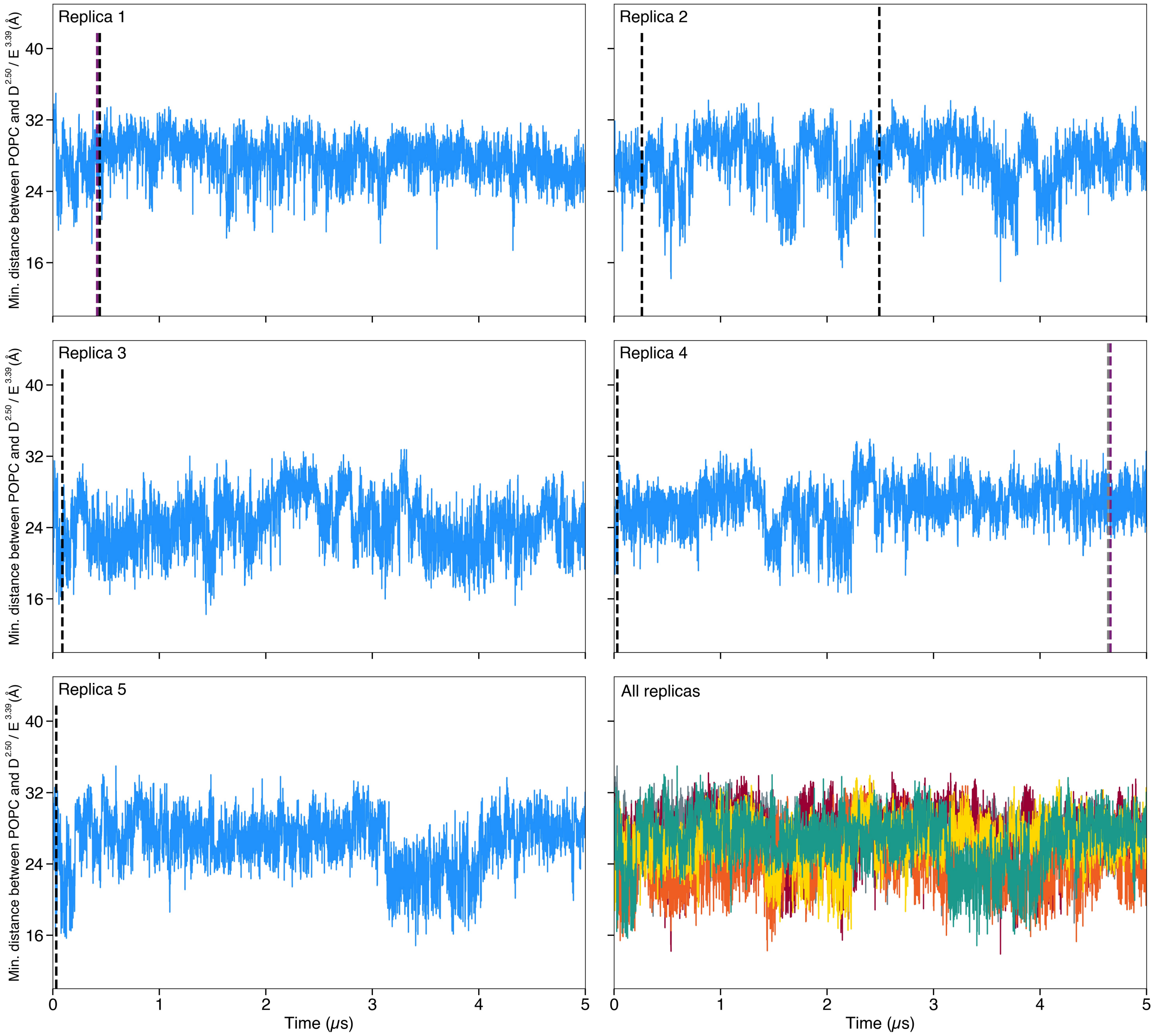


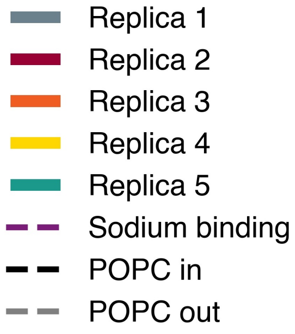


Figure S2: Time series of the sum of the minimum distance between the choline groups of POPC and residue D69$^{2.50}$ and the minimum distance between the choline groups of POPC and residue E110$^{3.39}$ in simulations without ions in the ion binding site. The vertical dashed lines indicate the time at which a given event was observed. Sodium binding to the ion binding site is depicted by purple dashed lines, while POPC snorkeling into and out of the ion binding site is represented by black and grey dashed lines, respectively. The bottom right graph displays the combined data from all five replicas.

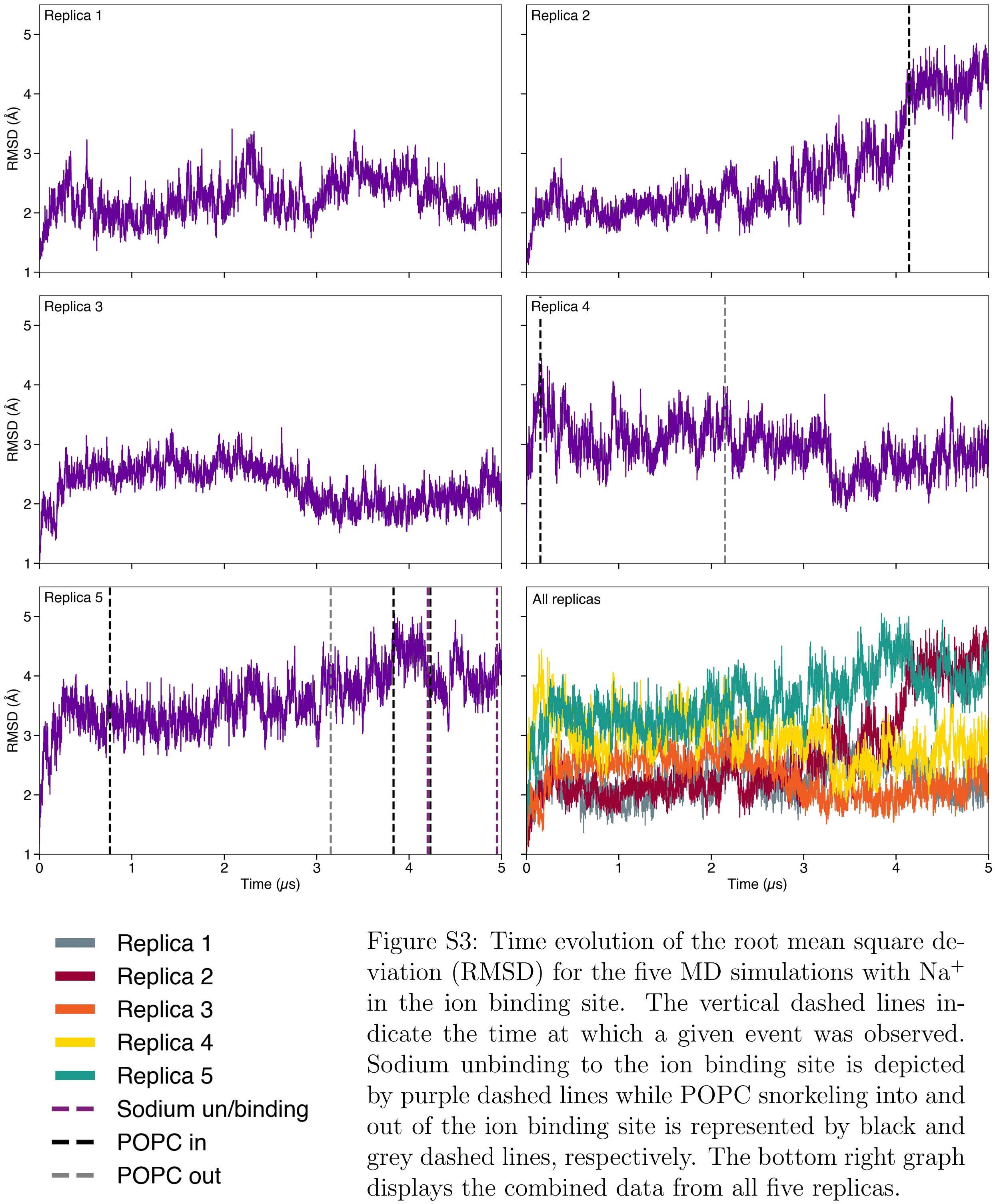


Figure S3: Time evolution of the root mean square deviation (RMSD) for the five MD simulations with $Na^+$ in the ion binding site. The vertical dashed lines indicate the time at which a given event was observed. Sodium unbinding to the ion binding site is depicted by purple dashed lines while POPC snorkeling into and out of the ion binding site is represented by black and grey dashed lines, respectively. The bottom right graph displays the combined data from all five replicas.

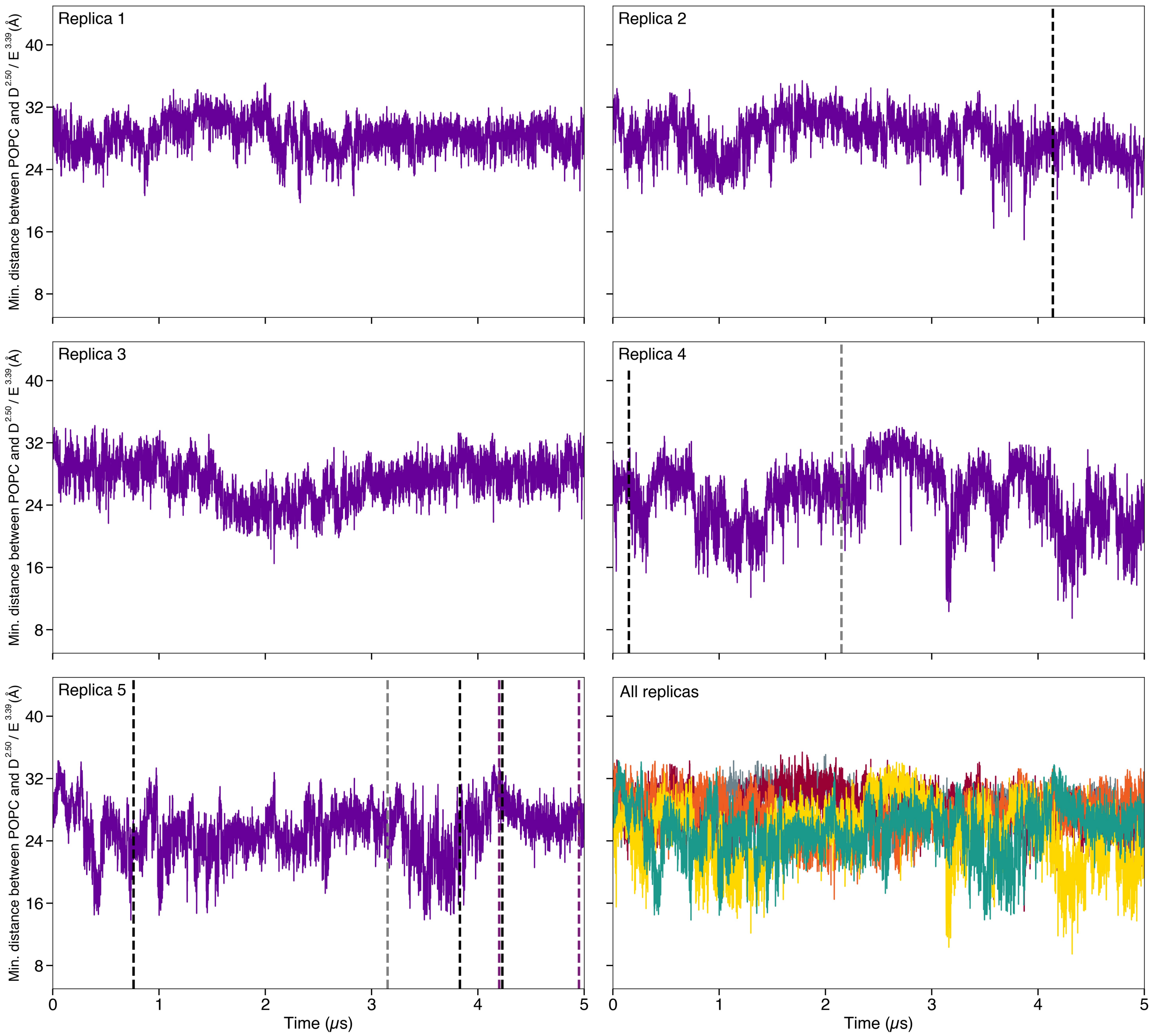


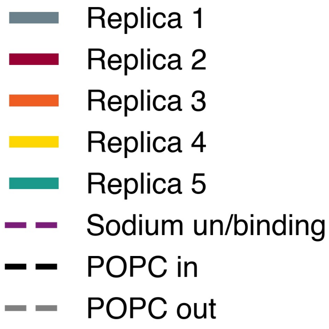


Figure S4: Time series of the sum of the minimum distance between the choline groups of POPC and residue D69$^{2.50}$ and the minimum distance between the choline groups of POPC and residue E110$^{3.39}$ in simulations with $Na^+$ in the ion binding site. The vertical dashed lines indicate the time at which a given event was observed. Sodium binding and unbinding in the ion binding site is depicted by purple dashed lines, while POPC snorkeling into and out of the ion binding site is represented by black and grey dashed lines, respectively. The bottom right graph displays the combined data from all five replicas.

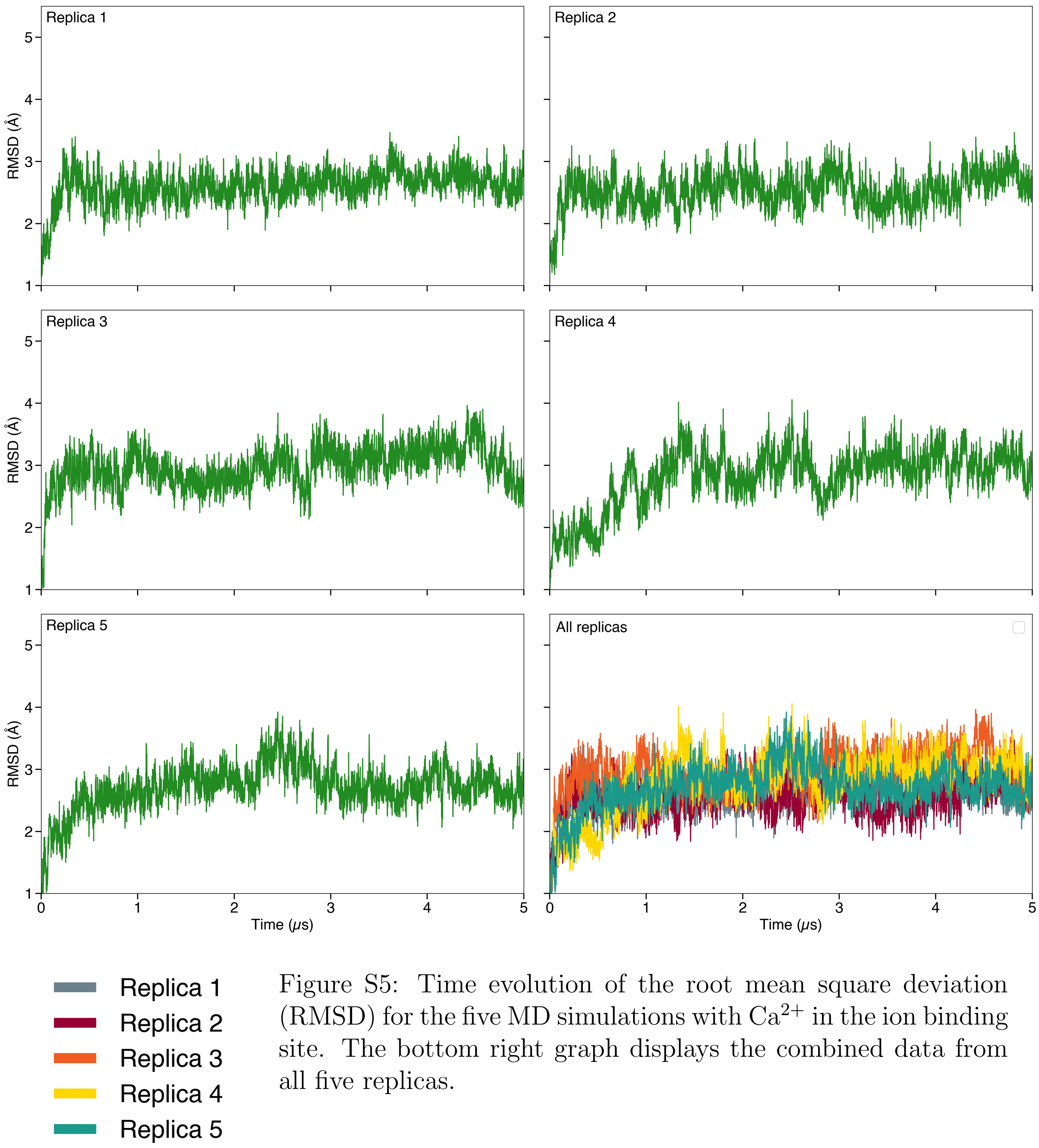


Figure S5: Time evolution of the root mean square deviation (RMSD) for the five MD simulations with $Ca^{2+}$ in the ion binding site. The bottom right graph displays the combined data from all five replicas.

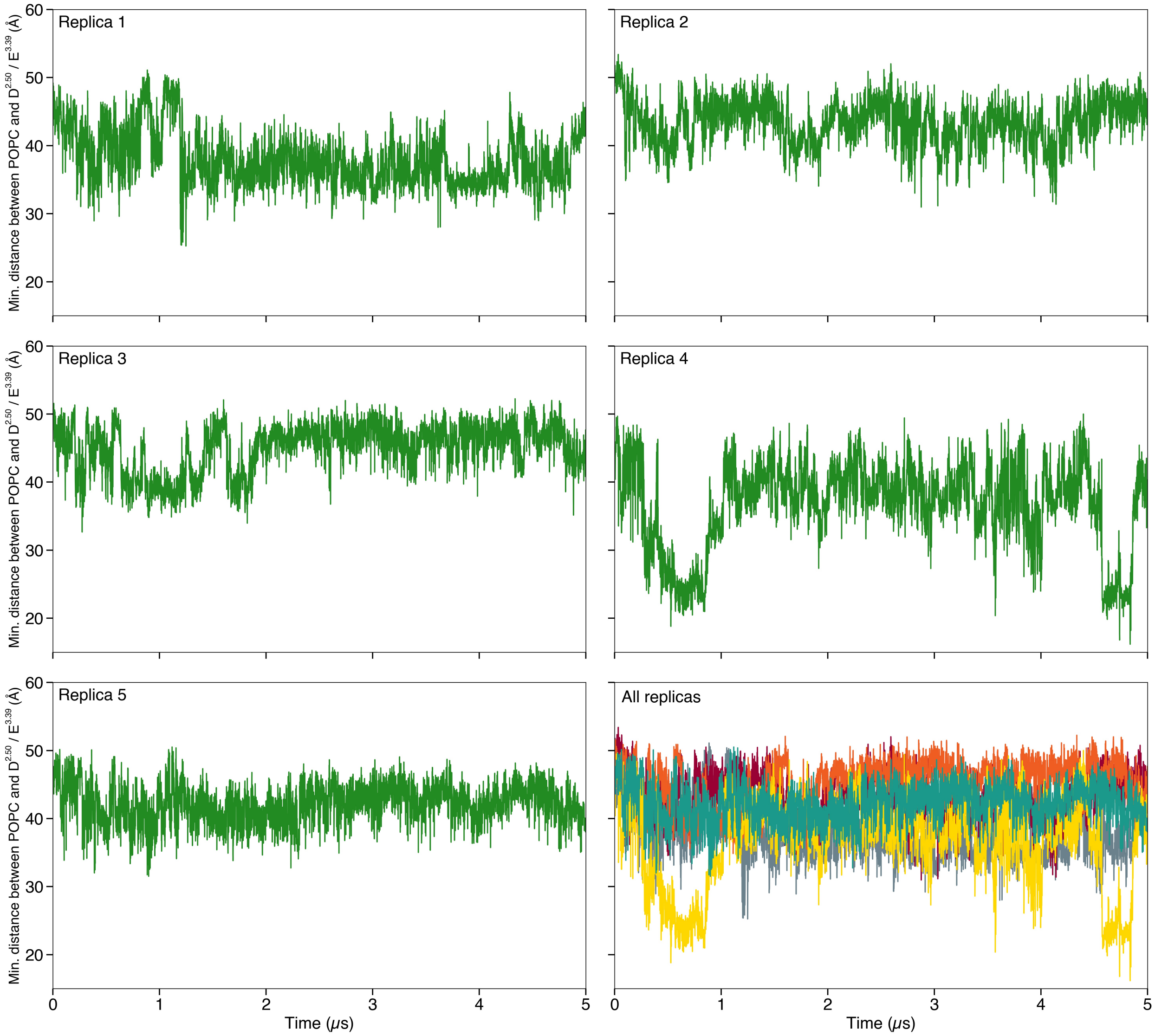


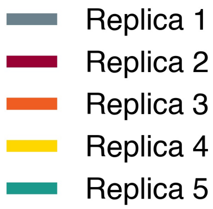


Figure S6: Time series of the sum of the minimum distance between the choline groups of POPC and residue D69$^{2.50}$ and the minimum distance between the choline groups of POPC and residue E110$^{3.39}$ in simulations with $Ca^{2+}$ in the ion binding site. The bottom right graph displays the combined data from all five replicas.

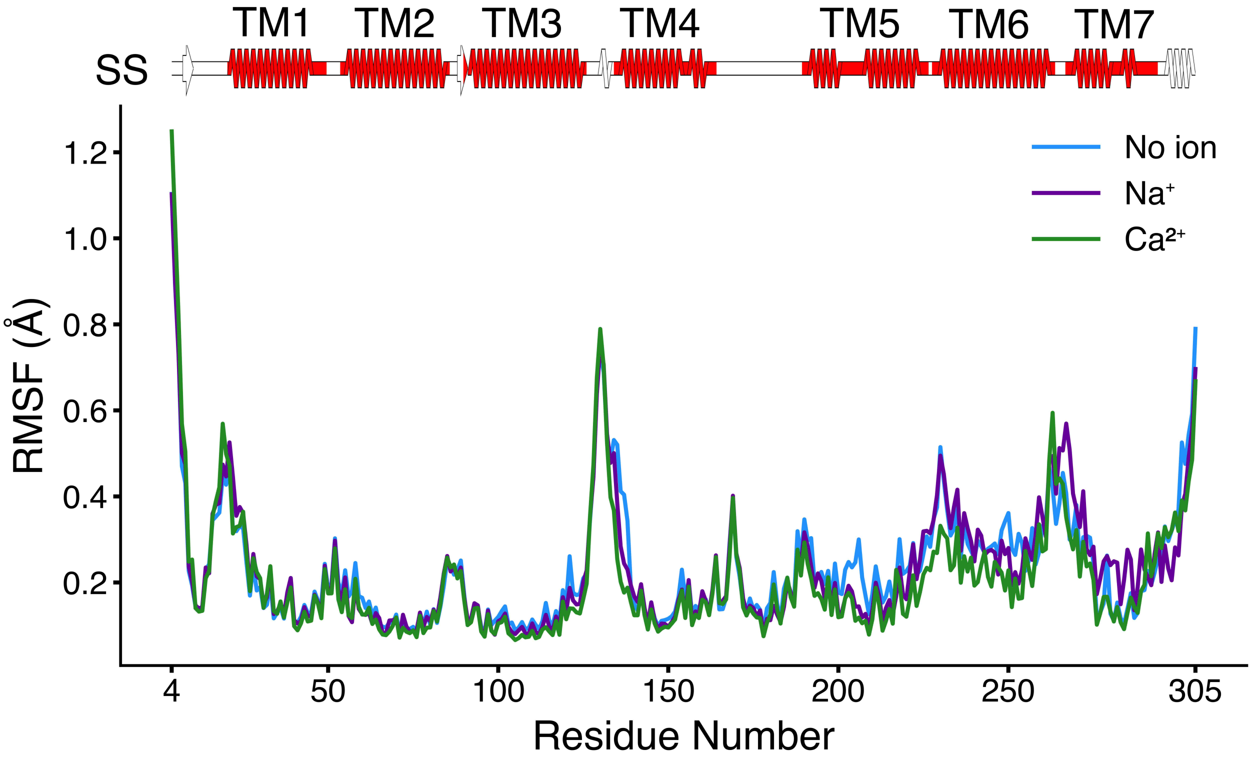


Figure S7: Root mean square fluctuation (RMSF) analysis with secondary structure representation. Top: secondary structure representation calculated and generated with SSDraw,[18] using as input the chain A of the OR51E2 cryo-EM structure (PDB ID: 8F76[1]). The transmembrane (TM) regions, as identified in table S1, are highlighted in red. Bottom: RMSF of the three sets of simulations.

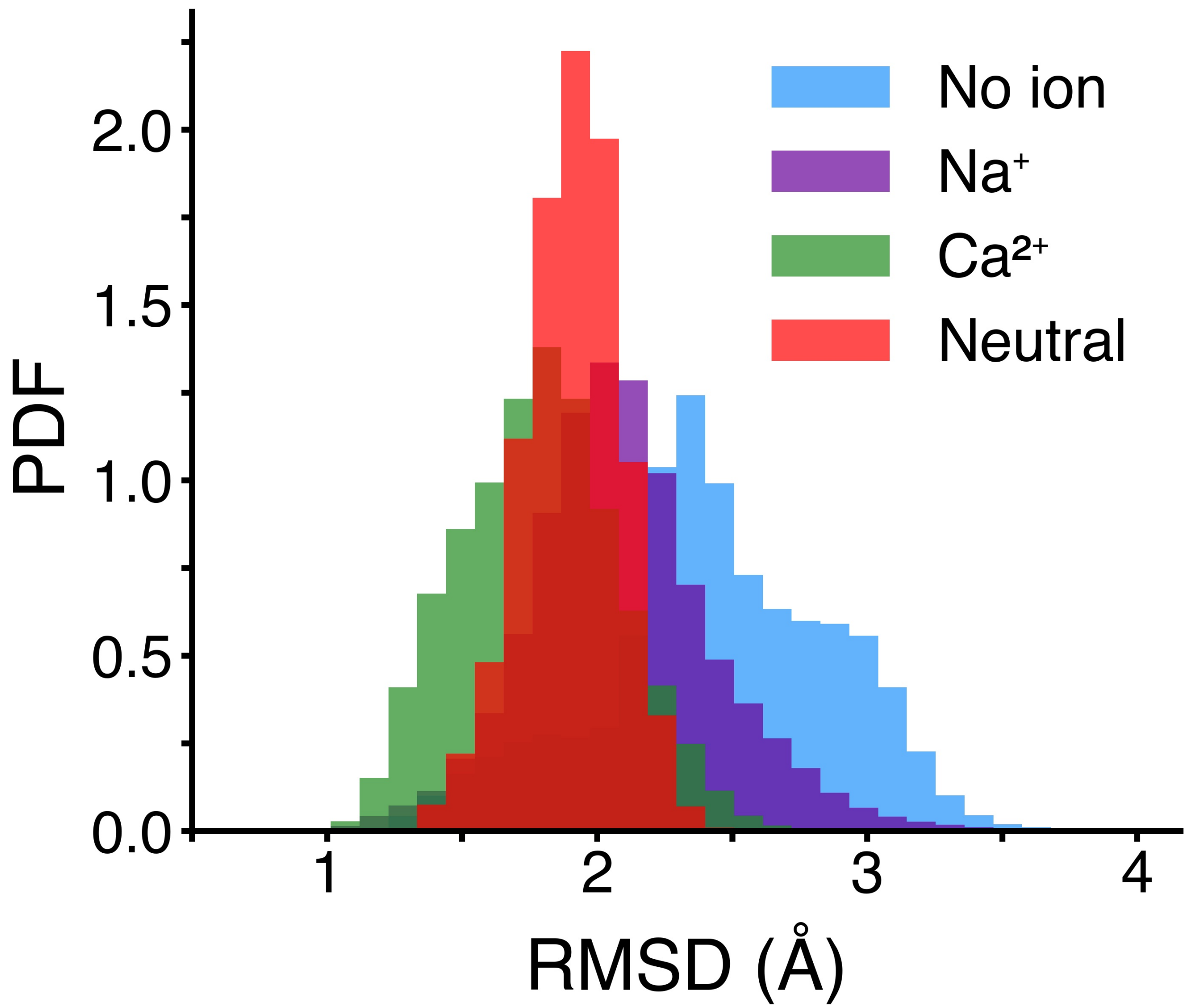


Figure S8: Histogram of the RMSD of the simulations with charged D69$^{2.50}$ and E110$^{3.39}$ performed in this work (without ions, with $Na^+$, and with $Ca^{2+}$ in the ion binding site) and of the simulations with neutral D69$^{2.50}$ and E110$^{3.39}$ with respect to TM3-TM5-TM6 $C_\alpha$ of the apo form of OR52$_{cs}$. We can observe that also quantitatively the simulations with neutral binding pocket sampled receptor conformations in between those of the $Na^+$ and $Ca^{2+}$ simulations.
